# Distinct cellular immune profiles in the airways and blood of critically ill patients with COVID-19

**DOI:** 10.1101/2020.10.29.360586

**Authors:** Anno Saris, Tom D.Y. Reijnders, Esther J. Nossent, Alex R. Schuurman, Jan Verhoeff, Saskia D. van Asten, Hetty J. Bontkes, Siebe G. Blok, Janwillem Duitman, Harm Jan Bogaard, Leo Heunks, Rene Lutter, Tom van der Poll, Juan J. Garcia Vallejo, on behalf of the ArtDECO consortium and the Amsterdam UMC COVID study group

**Author notes:** both authors contributed equally (Tom van der Poll, Juan J. Garcia Vallejo). **Corresponding author:** Anno Saris, *Center for Experimental and Molecular Medicine, Amsterdam UMC, Academic Medical Center, Amsterdam, The Netherlands.

## Abstract

Our understanding of the coronavirus disease-19 (COVID-19) immune response is almost exclusively derived from studies that examined blood. To gain insight in the pulmonary immune response we analysed BALF samples and paired blood samples from 17 severe COVID-19 patients. Macrophages and T cells were the most abundant cells in BALF. In the lungs, both CD4 and CD8 T cells were predominantly effector memory cells and expressed higher levels of the exhaustion marker PD-1 than in peripheral blood. Prolonged ICU stay associated with a reduced proportion of activated T cells in peripheral blood and even more so in BALF. T cell activation in blood, but not in BALF, was higher in fatal COVID-19 cases. Increased levels of inflammatory mediators were more pronounced in BALF than in plasma. In conclusion, the bronchoalveolar immune response in COVID-19 has a unique local profile that strongly differs from the immune profile in peripheral blood.

**Summary:** The bronchoalveolar immune response in severe COVID-19 strongly differs from the peripheral blood immune profile. Fatal COVID-19 associated with T cell activation blood, but not in BALF.

## Introduction

Severe acute respiratory syndrome coronavirus 2 (SARS-CoV-2) causes coronavirus disease 2019 (COVID-19) and is responsible for the current pandemic. At the time of writing, 21.6 million confirmed cases with over 767,000 deaths have been reported in 216 countries.^1^ The majority of infected individuals experiences no to mild symptoms, but 14% of infected persons develop severe and 5-6% critical life-threatening disease.^1,2^ From the hospitalized patients 20—30% require respiratory support in the intensive care unit (ICU), with an average of 6-8 weeks until clinical recovery,^2^ leading to an unprecedented strain on healthcare systems.^1^

Using the receptor binding domain (RBD) of the spike protein, SARS-CoV-2 infects host cells to ultimately induce cell death by pyroptosis,^3^ the most immunogenic form of cell death that induces a strong inflammatory response.^4^ The resulting wave of pro-inflammatory cytokines recruits other immune cells, mostly monocytes and T cells that act to clear the infection.^3,5,6^ It is hypothesized that both cytotoxic and humoral adaptive responses are necessary to efficiently control SARS-CoV-2 infection.^7^ Virus-specific T cells are observed in most patients,^8–10^ and we previously reported that the magnitude of antigen-specific T cell responses is unrelated to disease severity.^11^ In some patients, however, excessive release of cytokines is induced, for reasons that are currently unknown, giving rise to a cytokine storm that leads to severe lung damage and acute respiratory distress syndrome (ARDS).^3,5,6^ Ultimately, in 70% of fatal cases death is caused by respiratory failure due to ARDS, whereas 28% of fatal cases is due to sepsis-like cytokine storm associated multi-organ failure.^12^

Our understanding of the immune response to SARS-CoV-2 is limited by the fact that most studies until now have used plasma and blood cells. As characteristics of systemic immunity in ARDS differ strongly from responses in the bronchoalveolar compartment,^13^ these studies may fail to elucidate the main pathological feature of COVID-19: development of severe and progressive lung damage. Bronchoalveolar lavage fluid (BALF) of COVID-19 patients appears to be enriched in transcripts of CCL2/MCP-1 and CCL7, chemokines involved in recruiting inflammatory CCR2+ monocytes.^14^ Inflammatory monocyte-derived macrophages were the dominant cell type in the lungs during severe and critical COVID-19 in a study utilizing single-cell RNA sequencing.^15^ In severe COVID-19 patients, these inflammatory macrophages seemed less abundant, primarily due to clonal expansion of CD8 tissue-resident T cells (Trm). Using bulk RNA sequencing of blood and BALF mononuclear cells from COVID-19 patients, another study showed striking differences in expression of some of the tested genes in these different body compartments.^16^ However, these studies did not allow for definitive conclusions,^15,16^ and reports on immune cell profiling in the lungs of COVID-19 patients based on protein expression are lacking and our knowledge thus remains limited regarding immunity during COVID-19 at the site of infection. Consequently, an urgent need still exists to further our understanding of the pulmonary immune response in COVID-19.

The current study aimed to decipher the bronchoalveolar immune response during late-stage severe COVID-19 disease and to compare this with the systemic peripheral blood immune response. To this end, we isolated mononuclear cells from both BALF and blood of COVID-19 patients suffering from persistent ARDS. Combining spectral flow cytometry with measurements of soluble inflammatory mediators, we provide a comprehensive overview of the pulmonary and systemic immune response during late stage COVID-19.

## Results

### Patient population

Blood and BALF samples were obtained from 17 critically ill COVID-19 patients 1 to 31 days after a median ICU stay of 15 days [IQR: 9-19.5] (supplemental fig. 3A provides information on time of sampling and number of samples per patients). ICU mortality was 23.5% (4 of 17) and 90-day mortality 29.4% (5 of 17). Patient characteristics are listed in table 1. One patient used hydrocortisone prior to development of COVID-19 for adrenal insufficiency.

**Table 1:**
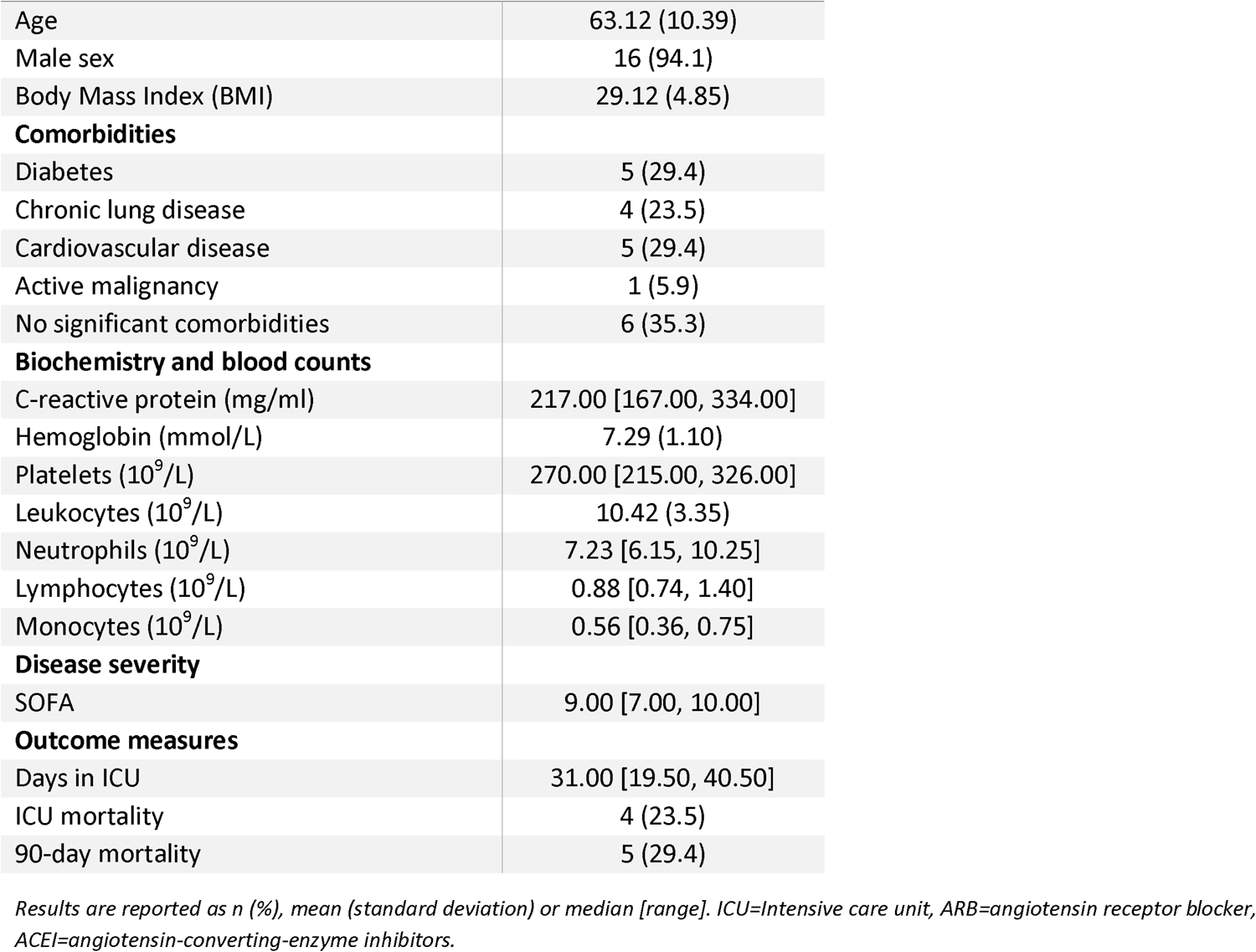
baseline characteristics and outcome (n=17)

### Immune phenotyping in blood and bronchoalveolar lavage

Paired PBMC and BALFMC samples were clustered unsupervised with overlaid colours of manual gates (Fig 1A-B) or staining intensity (supplemental figure 4+5). CD3+ T cells – taking into account their high abundance and plethora of subpopulations – were clustered separately throughout the manuscript to aid interpretation and facilitate more detailed analysis.

**Figure 1:**
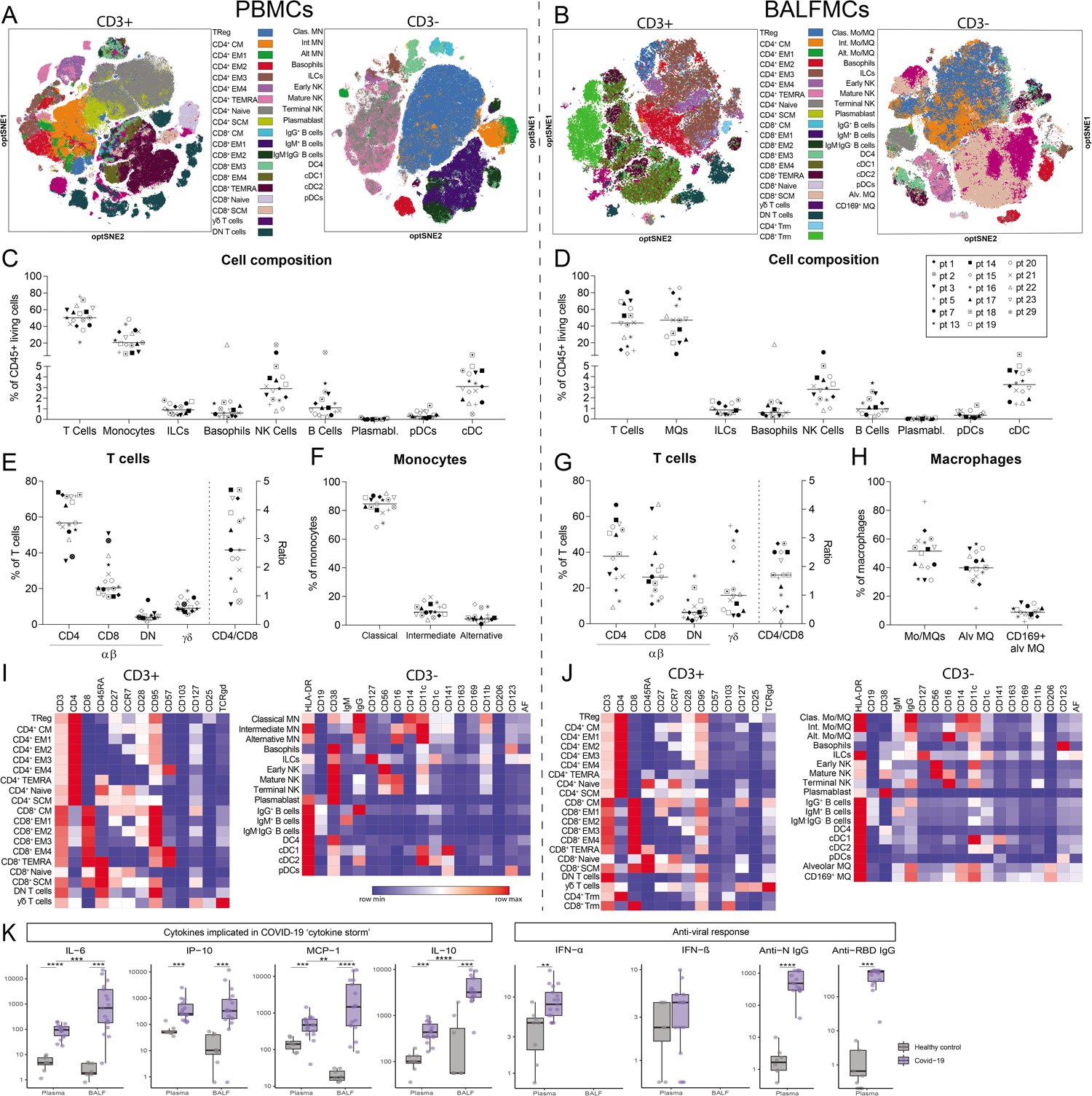
BALF predominantly comprises T cells and monocyte-derived and alveolar macrophages. PBMCs and BALFMCs isolated from COVID-19 patients admitted to the ICU were measured using spectral flow cytometry (n=17). (A+B) unsupervised analysis using omiq is presented in optSNE plots wherein colours are applied to clusters after manual gating (see supplementary figures 1+2). CD3+ and CD3-negative cells are depicted in separate optSNE plots. The dotted line separates data obtained from PBMCs and BALFMCs (C-H) quantification of general immune cell populations, T cell subsets and monocyte/macrophages subsets in PBMCs and BALFMCs. (I-J) Expression of lineage and exhaustion markers are depicted for all cell subsets using heat maps. ICU mortality was observed in patient 1, 3, 8 and 16. Statistical significance of cytokine levels was tested using Mann Whitney test (healthy vs COVID-19) or Wilcoxon (COVID-19 plasma vs BALF). *p<0.05, ** P<0.01, *** P<0.001, **** P<0.0001. Abbreviations: PBMC=peripheral blood mononuclear cells, BALFMC= bronchoalveolar lavage fluid mononuclear cells, Treg= regulatory T cells, CM= central memory T cells, EM= effector memory T cells, TEMRA=RA+ effector memory T cells, SCM=stem-cell like memory T cells, DN=double negative, Trm=tissue-resident memory T cell, MQ=macrophage, Mo/MQ=monocyte-like macrophage, Clas=classical, Int=intermediate, Alt=alternative, ILC=innate lymphoid cells, NK=natural killer cells, DC= dendritic cell, cDC=conventional DC, pDC=plasmacytoid DC, Alv=alveolar, αβ=αβ T cell receptor, ⍰δ=⍰δ T cell receptor.

In line with prior reports of lymphopenia in severe COVID-19 cases,^7,19^ peripheral blood lymphocyte counts were low: 0.88 [0.74, 1.40] x 10^6^/L (median, IQR) (Table 1). PBMC cell frequencies were within normal ranges,^20–23^ with limited variation between patients (Fig. 1C), but with CD4/CD8 ratios as high as 5 in some patients (normal range 1-3.6) (Fig. 1E). Cell composition in BALF, however, varied widely between patients. Macrophages and T cells were the most abundant and variable populations (46.7±25.0% and 42.5±23.9%; mean±SD; Fig. 1D). BALF CD4/CD8 ratios ranged from 0.1-2.8 and BALFMCs displayed remarkably high percentages of double-negative (DN) αβ T cells (8.5±6.8%) and ⍰δ T cells (22.2±17.7%; normal range 2-10%^24^) (Fig. 1G).

The expression of lineage, activation and exhaustion markers by the different cell populations is displayed in heat maps (Fig 1I-J). Compared to blood monocytes, monocyte-like cells in BALFMCs (further depicted as Mo/MQs) had a completely different expression of molecules such as CD38, CD1c, CD141 and CD11b. Fas receptor (CD95) was highly expressed on all, except naïve, T cell subsets in both PBMCs and BALFMCs. CD57, a marker for end-stage T cell exhaustion, was expressed in PBMCs on (terminally) differentiated EM3, EM4 and TEMRA CD4 and CD8 T cells, while CD57 expression was low in BALFMCs, which may suggest that TEMRA CD8 T cells in BALF are of the functional CD57-phenotype, while in circulation exhausted CD57+ TEMRA CD8 T cells were present.^25^

General inflammatory markers linked to COVID-19, (i.e. IL-6, C-X-C motif chemokine (CXCL)-10/IP-10, C-C motif ligand (CCL)-2/MCP-1),^3,5,6^ and also anti-viral IFN-α levels were significantly increased in both plasma and BALF of COVID-19 patients compared to healthy controls (Fig. 1K). Interestingly, levels of IL-6, CCL-2/MCP-1 and IL-10, but not CXCL10/IP-10, were higher in BALF than in plasma. All patients had developed an IgG antibody response targeting RBD of the spike protein and nucleocapsid protein (N) SARS-CoV-2 (Fig. 1K), the two dominant immunogenic proteins of SARS-CoV-2.^11^

### T cell differentiation and phenotypes in PBMCs and BALFMCs

Differentiation of T cells using CD95, CD28, CD27, CCR7 and CD45RA (see supplementary table 2 for details) revealed a high prevalence of central memory CD4 T cells and terminally differentiated CD8 T cells (TEMRA) in PBMCs (Fig. 2A+D). In BALFMCs however, (terminally differentiated) effector memory T cells were dominant wherein CD4 T cells were mainly EM2 and EM3 (Fig. 2A+D). In both plasma and BALF levels of interleukin (IL)-4, IL-17A and interferon (IFN)-0 were all below or around detection limit (Fig. 2I+J, *IFN-⍰ below detection limit),* precluding conclusions on T cell skewing. Granzyme B levels were highly upregulated in both plasma and BALF relative to values in control samples, and in patients BALF levels were higher than plasma levels (Fig. 2K). Effector CD8 T cells mainly had a Trm, EM4 and TEMRA phenotype. Based on CD38 and HLA-DR expression (gating strategy in supplemental Fig. 1+2), naïve and stem-like T cells did not show any activation *(data not shown),* while pronounced activation was observed in the other T cell subsets (Fig. 2B+E). Activated BALF T cells had an even higher PD-1 expression than PBMCs. Secretion of the PD-ligand 1 (soluble PD-L1) was significantly upregulated in both BALF and plasma of COVID-19 patients. IL-2 and IL-7, both stimulating T cell proliferation and differentiation,^21,27^ were upregulated in COVID-19 patients as compared to healthy control and IL-2 levels were higher in COVID-19 BALF as compared to COVID-19 plasma.

**Figure 2:**
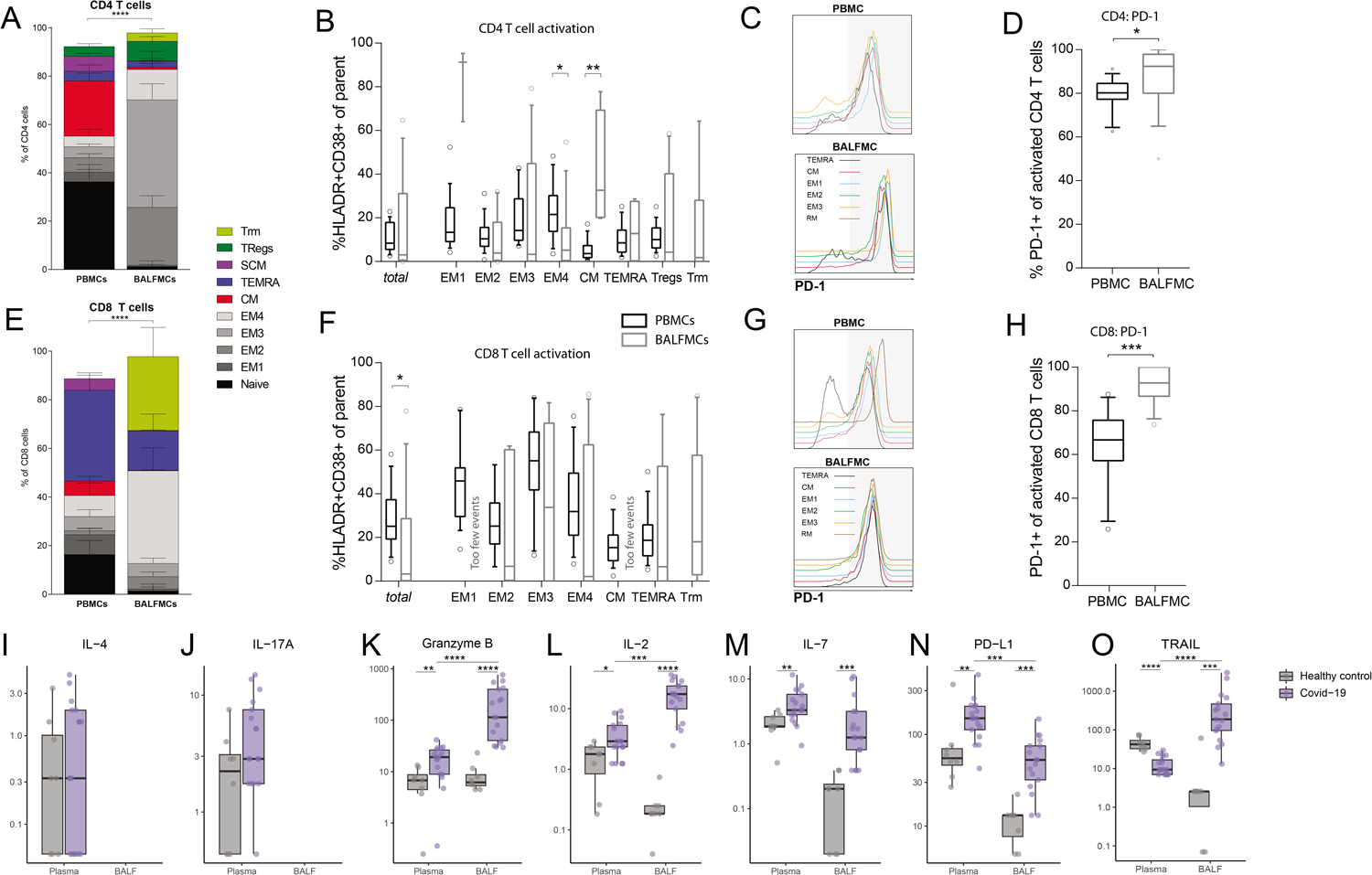
BALF T cells comprise predominantly effector memory CD4 and CD8 T cells and CD8 Trm. PBMCs and BALFMCs isolated from COVID-19 patients admitted to the ICU were measured using spectral flow cytometry (n=17). T cells were phenotyped using CD27, CD28, CCR7 and CD45RA and CD95 in naïve (CD45RA+CD27+CD28+CD95-), stemcell-like memory (SCM; CD45RA+CD27+CD28+CD95+), effector memory-1 (EM1) (CD45RA-CD27+CD28+CCR7-), EM2 (CD45RA-CD27-CD28+CCR7+), EM3 (CD45RA-CD27-CD28+CCR7-), EM4 (CD45RA-CD27-CD28-CCR7-), effector memory RA^+^ (TEMRA; CD45RA-CD27+CD28+CCR7-), central memory (CM; CD45RA-CD27+CD28+CCR7+), tissue-resident memory (Trm; CD103+CD28-; only for BALFMCs) and regulatory T cells (Treg; CD25++CD127-; only for CD4 T cells) (A+F). Activation (i.e. HLA-DR^+^CD38^+^) is presented for different CD4 and CD8 T cell subsets (only for populations with >250 events) (B+F). Representation of PD-1 expression on different T cells subsets (C+G) in PBMC and BALFMC with concomitant quantification of total PD-1 expression (D+H). Levels of IL-4 (I), IL17-a (J), Granzyme B (K), IL-2 (L), IL-7 (M), IL-10 (N) and soluble PD-L1 (O) are presented in plasma and BALF. Box plots represent median±interquartile range. Abbreviations: PBMC=peripheral blood mononuclear cells, BALF= bronchoalveolar lavage fluid, DN=double negative, PD-1=programmad death-1, PD-L1= programmed death-ligand 1, Anti-N=anti-nucleacapsid, Anti-RBD=anti-receptor binding domain of spike protein. Statistical significance was tested with Kruskal-Wallis (A+C+E+G), Wilcoxon (D+H, I-O: COVID-19 plasma vs BALF) or Mann Whitney test (I-O: healthy vs COVID-19). * p<0.05, ** P<0.01, *** P<0.001, **** P<0.0001

### Correlation PBMC and BALF population

Lymphopenia is a frequently reported phenomenon in COVID-19, which may either be caused by massive T cell migration into the lungs or activation-induced apoptosis.^3,7,28^ If lymphopenia would be caused by massive migration of a specific T cell subset, relative T cell numbers and CD4/CD8 ratios in PBMCs should inversely correlated with BALFMCs. However, relative T cell numbers in blood and BALF did not show any correlation (rho= −0.01, p=0.96), while CD4/CD8 ratios correlated significantly between blood and BALF (rho=0.65, p=0.0071) (Fig. 3A). CD4/CD8 ratios were lower in BALF, which shows a relative higher abundance of CD8 T cells in BALF. To gain insight in immune cell migration and the peripheral immune system versus pulmonary, PBMC subsets were correlated to BALFMC subsets (Fig. 3B). When comparing PBMCs to BALFMCs, the activation status of peripheral CM CD4 T cells correlated positively with conventional T cells and negatively with mo/MQs in BALF (Fig. 3C), which may be caused by activation induced proliferation of CM CD4 T and subsequent migration, causing higher T cell numbers in BALF (and consequently other cells will be less abundant). Furthermore, peripheral Tregs negatively correlated with DN T cells in BALF, which may indicate impaired induction of Fas-mediated T cells apoptosis by Tregs what has been shown to cause accumulation of DN T cells.^29^

**Figure 3:**
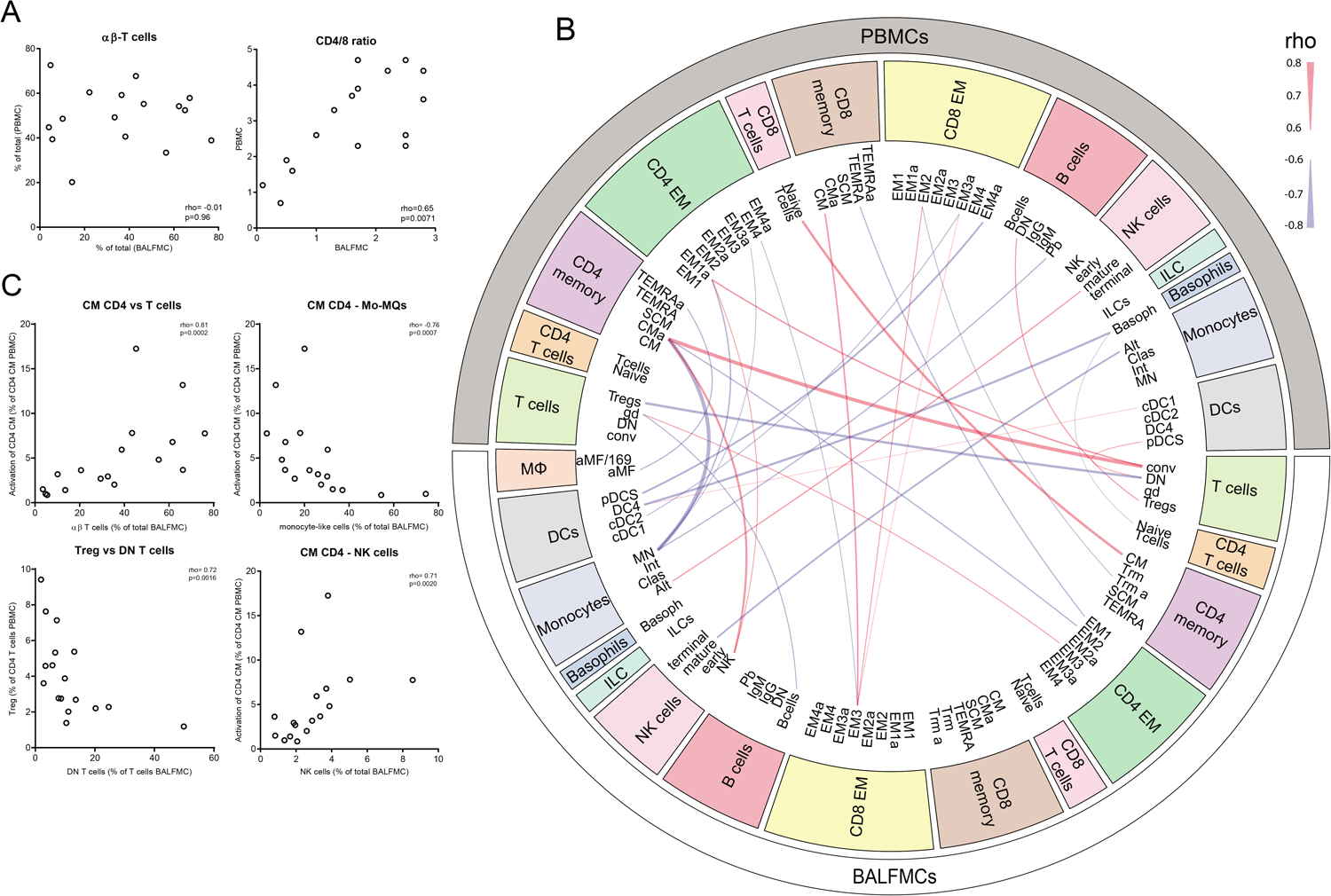
Correlations between cell populations in PBMCs and BALFMCs. The bronchoalveolar and systemic immune response were compared by correlating αβ-T cells and CD4/CD8 in PBMC and BALFMC (A) and comparing plasma and BALF cytokine levels (B). Cytokines are ordered in ‘implicated in COVID-19 cytokine storm’ (IL-6, CXCL10/IP-10 and CCL2/MCP-1) and ‘anti-viral responses’ (IFN-α, IFN-β, anti-RBD IgG and anti-N IgG) with box plots displaying median±interquartile range. All PBMC populations obtained using manual gating were correlated to BALFMC populations using spearman correlation. All significant correlations with a rho>0.6 are depicted in a circus plot wherein red depicts a positive correlation and blue depicts a negative correlation and line thickness resembles the goodness-of-fit (i.e. rho) (C). The four populations with the strongest correlation are presented in dot plots (D). Abbreviations: PBMC=peripheral blood mononuclear cells, BALFMC= bronchoalveolar lavage fluid mononuclear cells, Treg= regulatory T cells, CM= central memory T cells, EM= effector memory T cells, TEMRA=RA+ effoctor memory T cells, SCM=stem-cell like memory T cells, DN=double negative, Trm=tissue-resident memory T cell, MQ=macrophage, Mo/MQ=monocyte-like macrophage, Clas=classical, Int=intermediate, Alt=alternative, ILC=innate lymphoid cells, NK=natural killer cells, DC= dendritic cell, cDC=conventional DC, pDC=plasmacytoid DC, Alv=alveolar, αβ=αβ T cell receptor, ⍰δ=⍰δ T cell receptor.

### Influence of duration of ICU stay on bronchoalveolar and systemic immune responses

To investigate the change in cellular composition of the immune system during prolonged ICU stay, samples were grouped according to ICU stay with a cut-off of 14 days (<14 days: n=10, 9 days [8-12.5]; >14 days: n=9, 18 days [17-22.5]) (median [interquartile range]) (see supplemental fig. 3B). We included only one sample per patient in each group. Unsupervised clusters clearly showed a significant effect of ICU stay on immune cell composition, especially in T cells in BALF (Fig. 4). The differences in BALF T cells mainly resulted from lower activation of multiple T cell subsets in the ICU stay > 14 days group: CD4 EM2 (1.4±0.7% vs 14.3±4.5% of CD4 EM2), CD4 EM3 (2.7±1.8% vs 39.1±9.6% of CD4 EM3), CD8 EM4 (3.8±3.2% vs 47.3±11.6% of CD8 EM4), CD8 TEMRA (2.8±2.6% vs 27.7±9.8% of CD8 TEMRA) (Fig. 4D). Furthermore, conventional (i.e. αβ) T cells were less abundant (50.0±7.1% vs 21.5±6.6%), whereas the frequencies of alveolar macrophages, monocyte-like macrophages and ⍰δ T cells tended to be higher after prolonged ICU stay. PBMCs isolated after >14 days ICU stay also exhibited an overall trend towards lower frequencies of activated T cell subsets, but the differences were less pronounced than in BALFMCs. Plasma levels of CXCL10/IP-10 were lower with extended ICU stay, while in BALF ICU stay only affected TNF-related apoptosis-inducing ligand (TRAIL) levels. IL-6, CCL-2/MCP-1, IL-2, PD-L1 and granzyme B remained stable during extended ICU stays.

**Figure 4:**
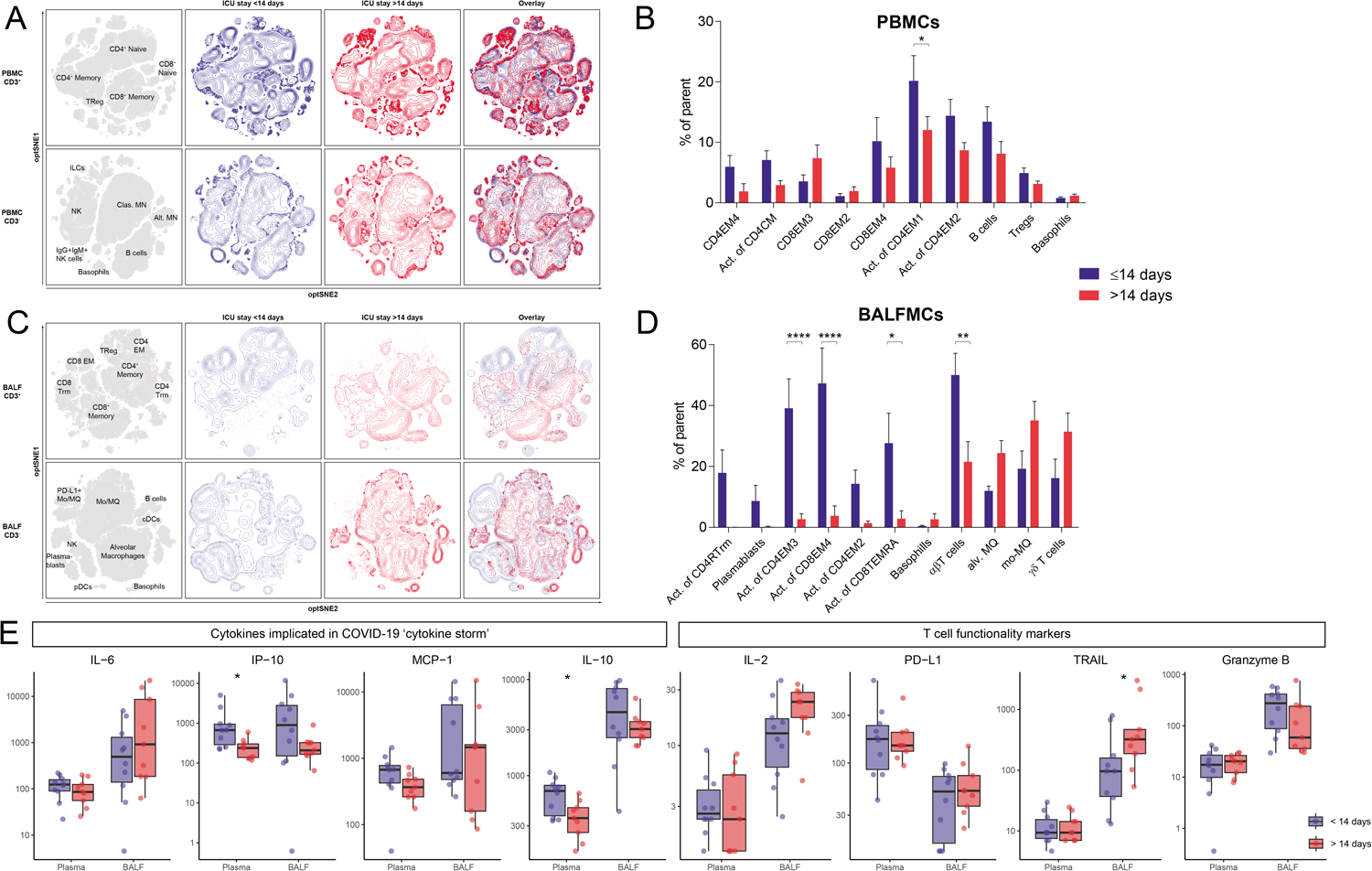
Influence of duration of ICU stay on bronchoalveolar and systemic immune responses. Samples were stratified based on moment of sampling in ≤14 days (n=8) and >14 days (n=9). Only one sample per patient was included in each group. Immune cell population from PBMCs (A) and BALFMCs (C) were clustered using omiq unsupervised clustering and presented as optSNE plots. The ten populations with biggest relative differences (sorted from left to right) are depicted for PBMC (B) and BALFMC (D). Cytokine levels in plasma and BALF, as measured using luminex, were compared ≤14 days and >14 days of ICU stay wherein box plots represent median±interquartile range (E). Abbreviations: PBMC=peripheral blood mononuclear cells, BALFMC= bronchoalveolar lavage fluid mononuclear cells, Treg= regulatory T cells, CM= central memory T cells, EM= effector memory T cells, TEMRA=RA+ effoctor memory T cells, Trm=tissue-resident memory T cell, MN=monocyte, Mo/MQ=monocyte-like macrophage, ILC=innate lymphoid cells, NK=natural killer cells, DC= dendritic cell, cDC=conventional DC, pDC=plasmacytoid DC, Alv=alveolar. Statistical significance was tested using two-way ANOVA with correction for multiple testing using Holm-Sidak (B+D) or Mann Whitney test (E). * p<0.05, ** P<0.01, *** P<0.001, **** P<0.0001

### Association of fatal COVID-19 with bronchoalveolar and systemic immune responses

To dissect which immunological profile associates with mortality, patients were stratified according to survival (13 survivors and 4 non-survivors). Peripheral lymphocyte counts did not differ between survivors (0.85 [0.74, 1.31] x 10^6^/L) and non-survivors (1.53 [0.69, 2.77] x 10^6^/L, p=0.40). Samples were obtained 15.5 [12.5-17] days or 16 [9-22) days (median [interquartile range]) after ICU admission, respectively (see supplemental fig. 3C). Clustering surviving and deceased patients separately revealed large differences in both CD3- and CD3+ unsupervised clusters of BALFMCs (Fig 5A+D). Activation of different T cell subsets in blood was increased in non-surviving patients as compared to surviving patients: CD4 EM4 (32.7±4.3% vs 16.1±2.9% of CD4 EM4), CD4 EM3 (35.3±6.0% vs 18.2±2.5% of CD4 EM3) and CD8 TEMRA (31.2±6.9% vs 17.3±2.6% of CD8 TEMRA) (Fig. 5B). In contrast, activation of different T cells subsets in BALF tended to be reduced in non-surviving patients as compared to surviving patients: CD8 TEMRA (2.9±2.8% of CD8 TEMRA vs 17.3±7.6% of CD8 TEMRA), CD4 Trm (3.5±3.5% vs 11.8±6.0% of CD4 Trm), CD8 EM4 (12.3±11.7% vs 27.3±10.3% in CD8 EM4), and CD4 EM3 (12.2±10.9% vs 23.3±8.3% of CD4 EM3) (Fig. 5D). Although hampered by low sample size, the frequency of plasmablasts and ⍰δ T cells tended to be reduced in BALF of fatal cases as compared to surviving cases, while basophils were high in some fatal cases (5.1±3.9% vs 0.6±0.1%), which may be interesting to study further given the strong IgA response,^30^ and profound complement activation in COVID-19,^29^ both capable of stimulating histamine release.^31,32^ The levels IL-6, CXCL10/IP-10, CCL2/MCP-1, anti-SARS-CoV-2 IgG antibodies, granzyme B, IL-2 and TRAIL were not significantly different in both plasma and BALF of non-surviving compared to surviving patients.

**Figure 5:**
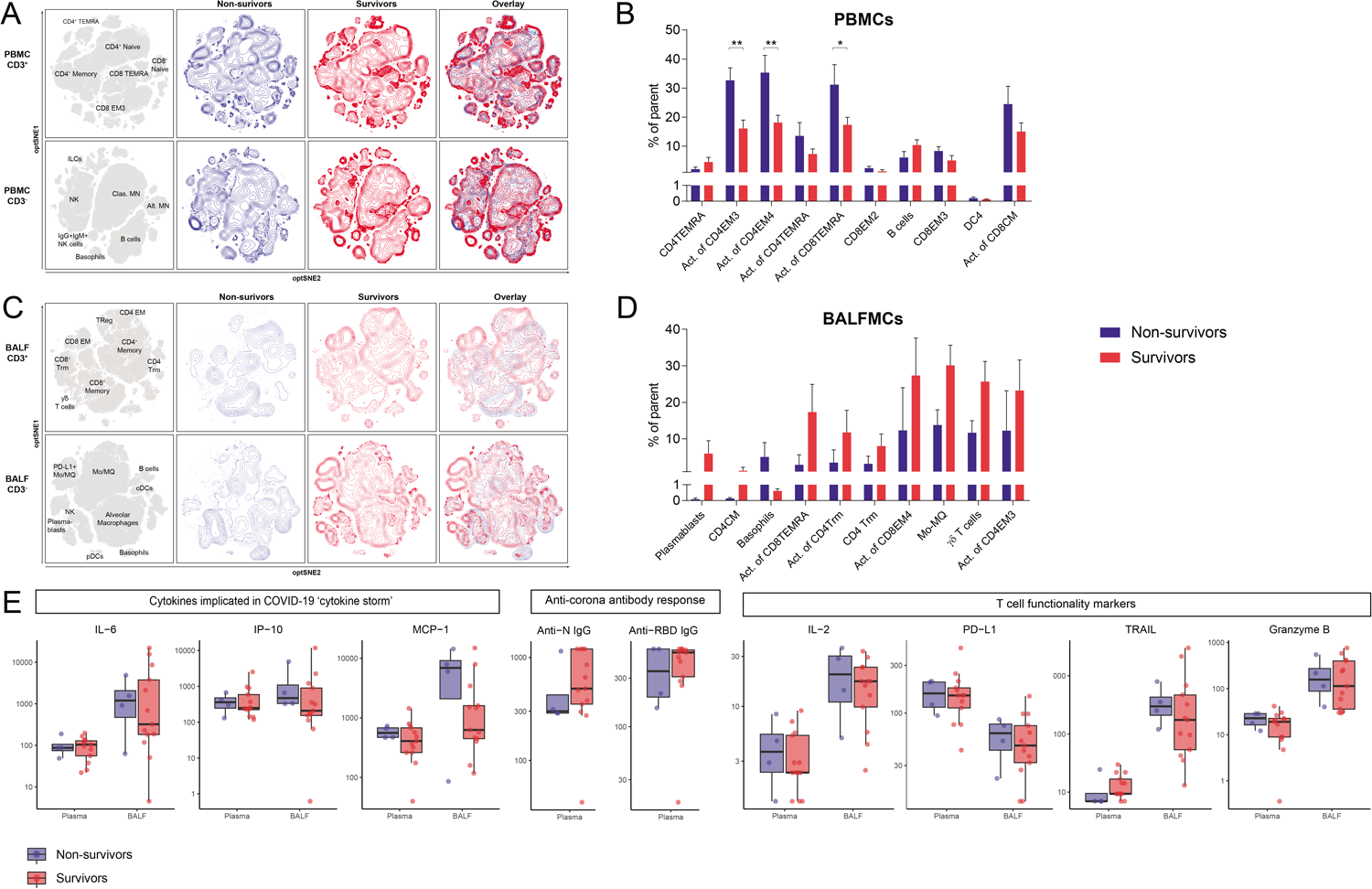
Association of fatal COVID-19 with bronchoalveolar and systemic immune responses. Samples were stratified based on ICU mortality (n=4 non-survivors (blue) vs n=13 survivors (red)). Only one samples per patient was included in each group N=. (A+C) Immune cell population from PBMCs (A) and BALFMCs (C) were clustered using omiq unsupervised clustering and presented as optSNE plots. (B+D) The ten populations with biggest relative differences (sorted from left to right) are depicted for PBMC (B) and BALFMC (D). Cytokine and antibody levels in plasma and BALF, as measured using Luminex and ELISA respectively, were compared in fatal and non-fatal COVID-19 cases and presented in box plots as median±interquartile range. (E). Abbreviations: PBMC=peripheral blood mononuclear cells, BALFMC= bronchoalveolar lavage fluid mononuclear cells, Treg= regulatory T cells, CM= central memory T cells, EM= effector memory T cells, TEMRA=RA+ effector memory T cells, Trm=tissue-resident memory T cell, MN=monocyte, Mo/MQ=monocyte-like macrophage, ILC=innate lymphoid cells, NK=natural killer cells, DC= dendritic cell, cDC=conventional DC, pDC=plasmacytoid DC. Statistical significance was tested using two-way ANOVA with correction for multiple testing using Holm-Sidak (B+D) or Mann Whitney test (E). * p<0.05, ** P<0.01, *** P<0.001, **** P<0.0001

## Discussion

Although a tremendous global effort of the scientific community has greatly improved our understanding of COVID-19, the exact pathophysiology remains to be unravelled. Some studies have reported overactivation of the immune system, whereas others have shown T cell exhaustion, dysfunction and/or apoptosis.^3,7,8,10,33^ Importantly, however, these earlier investigations were restricted to immune responses in blood. The present study shows a highly diverse immune response in COVID-19 patients admitted to the ICU, with significant differences between immune cells isolated from blood and BALF. The strongest correlation between PBMCs and BALFMCs was the activation status of peripheral CM CD4 T cells, which positively correlated with conventional T cells and negatively with mo/MQs in BALF. In COVID-19 patients, proportions of macrophages and T cells were highly variable in BALF. When comparing BALF with blood, T cell differentiation was highly skewed to an effector memory phenotype with a significantly higher PD-1 expression. These results indicate that obtaining cells from the lungs is pivotal in order to fully understand COVID-19 pathophysiology.

Despite previous reports of significant lymphopenia in COVID-19 patients upon admission,^3,7,28,34^ lymphopenia of ICU-admitted COVID-19 patients was only mild in our study. This may have resulted from late sampling, as lymphocyte numbers are reported to recover in a subset of patients.^34^ Severe lymphopenia is hypothesized to be caused by activation-induced cell death or massive migration of T cells to the site of infection.^7^ Here we show that relative peripheral T cell numbers do not inversely correlate with relative T cell numbers in BALF, which renders massive migration of T cells into the lungs an unlikely explanation for lymphopenia in prolonged critically ill COVID-19 patients. Fas expression, which has previously been correlated to the extent of peripheral lymphopenia,^5^ was high on all T cells in blood, except naïve T cells, which suggests that apoptosis is a more likely cause of T cell lymphopenia in COVID-19. In line with previous reports,^7,8^ PD-1 expression on activated peripheral blood T cells was high in our study, but it was even higher in BALFMCs, reaching over 95% positivity in most samples. This, combined with a lower Fas expression, may indicate that T cells in the bronchoalveolar space are more exhausted than those in blood, but less vulnerable to Fas-mediated apoptosis.

Tissue-resident CD8 T cells (CD8 Trm) were recently associated with moderate disease, while the inflammatory monocyte-like macrophages (mo/MQs) were more abundant in critical cases.^15^ In the present study, however, we did not observe a positive correlation of CD8 Trm with survival. If anything, mo/MQs were lower in fatal COVID-19. There are multiple possible explanations for these diverging observations: low sample sizes, different clinical comparisons (i.e. disease severity in all COVID-19 patients *versus* mortality in critical cases only), and time of sampling relative to disease onset (4-10 days after hospital admission^15^ compared to 1-30 days after ICU admission).

During COVID-19 a significant differentiation of peripheral blood CD4 and CD8 T cells into an effector memory phenotype has been reported,^35^ but little is known about T cell phenotypes at the primary site of infection, i.e. the lungs. The present study shows that differentiation in BALF is even more skewed, with a striking 89.6% of CD4 and 96.1% of CD8 T cells in BALF having effector memory phenotypes. T cell activation in both blood and BALF diminished with prolonged ICU stay (i.e. >14 days), which is consistent with suggestions of exhausted T cells in COVID-19. Remarkably however, while activation of T cells in the lungs seemed reduced in fatal COVID-19, peripheral T cell activation was increased, including activation of CD4 EM3 cells. These cells have a high cytolytic activity with corresponding high granzyme B production,^36^ and granzyme B was increased in both BALF and plasma when compared to uninfected controls. Our results show that both T cell activation and PD-1 expression on T cells are increased during COVID-19. Increased T cell activation in peripheral circulation was associated with mortality, with no evidence of activation in the lungs. The contrast in T cell activation in surviving *versus* non-surviving patients could theoretically result from a systemic activation of T cells, or a failure of activated T cells to migrate into the lungs in fatal COVID-19 cases, but this requires further investigation.

By investigating inflammatory markers as well as mononuclear immune cells in both peripheral blood and the lungs, our study provides an extensive overview of the immune response in late-stage critically COVID-19 patients. More importantly, this in-depth comparison of the peripheral blood and pulmonary compartments revealed stark differences in immune responses (e.g. T cell activation) that must be taken into account when using peripheral blood cells as a surrogate for the entire COVID-19 immune response. Moreover, while previous investigations have reported on a so-called systemic cytokine-storm in COVID-19,^3,5,6^ our study shows that the BALF levels of many cytokines are higher than their plasma levels (in spite of the dilution caused by the BAL procedure), indicating that a local rather than systemic cytokine storm is at play during late stage COVID-19. The present study is limited by its relatively low sample size – due to a rapidly declining incidence of SARS-CoV-2 infections after governmental measures were introduced – and variation in time of sampling, and these limitations necessitate caution when drawing conclusions.

In conclusion, immune composition in the lungs of COVID-19 patients admitted to the ICU was substantially different than in peripheral blood. BALF mainly comprised macrophages and T cells, with high percentages of inflammatory monocyte-like macrophages and ⍰δ T cells especially after prolonged ICU stay. Both CD4 and CD8 T cells expressed higher levels of PD-1 in BALF as compared with PBMCs. Surprisingly, total CD8 T cell activation in BALF was lower than in peripheral blood. Reduced CD4 and CD8 T cell activation associated with extended ICU stay, especially in BALF, while peripheral activation of T cells (CD4 EM3 and EM4 as well as CD8 TEMRA) associated with mortality.

## Materials and methods

### Subjects and study approval

The current study was part of the Amsterdam Study for DEep Phenotyping of COVID-19 disease (ArtDECO) 1 study, a cohort study of COVID-19 patients with persistent ARDS (mechanical ventilation > seven days). Per clinical protocol obtained left over biological samples were stored in the anonymized research Amsterdam UMC COVID-19 biobank (#2020-182). Informed consent for the use of samples and data was deferred until discharge from the ICU. In case of death, informed consent was requested from the patient’s relatives. Study procedure was approved by the Review Committee Biobank of the Amsterdam UMC (2020-065). All patients from whom BALF mononuclear cells were available were included in the present study. The study is in accordance with the declaration of Helsinki and adheres to Dutch regulations. Healthy controls were recruited as age-matched controls for two explorative studies, the explorative RILCA and RILCO trials (NL48912.018.14 and NL53354.018.15). Healthy controls had no (history of) respiratory disease or comorbidities and were 18-50 (RILCA) and 40-70 years (RILCO) of age, non-allergic, non-smoking or ex-smokers for at least one year and had a BMI of 17-30 kg.m_2_. Both the RILCA and RILCO studies were approved by the internal IRB and participants provided written informed consent.

### Isolation of BALF, plasma, peripheral blood mononuclear cells and BALF mononuclear cells

Prior to diagnostic bronchoalveolar lavage in COVID-19 patients, venous blood was drawn in EDTA and heparin tubes. EDTA blood was centrifuged 10 min at 1800g and supernatant plasma was collected and stored at −80°C. Peripheral blood mononuclear cells (PBMCs) were isolated from heparinized blood samples using standard Ficoll-Paque density gradient centrifugation (1000g, 20min, 21°C). During diagnostic bronchoscopy 2 x 20 ml 0,9 % NaCl at a (sub)segmental level, each aspirated immediately with low suction. From this, 10ml was used for microbiological diagnostic purposes and the remaining (~3-20ml) was centrifuged (300g, 10min, 4°C). BALF supernatant was stored at −80° and cell pellet was resuspended in 2mM dithiotreitol (Sigma, Zwijndrecht, the Netherlands) to solubilize sputum and mucus. After 30 min at 4°C, cells were washed with PBS+1%BSA and mononuclear cells were isolated using Ficoll-Paque density gradient centrifugation. PBMCs and BALF mononuclear cells (BALFMCs) were cryopreserved in liquid nitrogen until further analysis. Healthy BALF was collected by instilling eight successive 20 ml aliquots of pre-warmed 0.9% NaCl instilled at a (sub)segmental level, each aspirated immediately with low suction, according to the recommendations of the NHLBI and the National Institute of Allergy and Infectious Diseases.^17^ Fractions 3-8 were pooled, centrifuged for 10 minutes at 267g at 4°C and the supernatant was stored at −80°C till further analyses.

### Spectral flow cytometry

Cells were thawed and subsequently washed in IMDM+10%FCS+75U/ml DNAse (Sigma). To minimize day-to-day variation, all samples were thawed and stained on the same day. Next, 1 million cells (or as many as available in case of BALFMCs) were stained with live/dead stain, and subsequently monoclonal antibodies added sequentially: CCR7, ⍰δ T cell receptor, all non-brilliant (ultra) violet (BV/BUV) or non-brilliant blue (BB) labelled markers and finally all BV, BUV and BB labelled antibodies. See supplemental table 1 for a list of all antibodies used for this staining. Cells were measured using a 5 laser Aurora system (Cytek Biosciences, Fremont, CA) and data analysed with SpectroFlo (Cytek Biosciences, Fremont, CA) and OMIQ (OMIQ Inc, Santa Clara, CA). Six of thirty-one BALF samples were excluded because too few viable CD45 + cells were measured (i.e. less than 2.000) and another five samples were excluded due to active prednisolone therapy as part of standard clinical care. PBMC samples were only included if a paired BALF sample was available. CD3+ and CD3-cells were analysed separately in each compartment and to cater for the different yield of viable CD45+ cells in cluster analyses, a maximum of 45.000 PBMC (both CD3+ or CD3-) and 10.000 CD3+ and 30.000 CD3-BALFMC were included from each sample. Manual gating strategy is depicted in supplemental figure 1 (PBMCs) and 2 (BALFMCs)

### Measurements of soluble immunological mediators

Cytokines and chemokines were measured using Human Magnetic Luminex Assay according to the manufacturer’s instructions (#LKTM014, R&D systems, MN, USA). Samples that were above the upper limit of quantification were set at the upper detection limit of the assay, while samples below detection limit were set at half of the of the lower detection limit. Anti-RBD and anti-NP IgG antibodies were measured in EDTA plasma samples at 100-1200 fold dilutions using ELISA as previously described.^11^ Plates were coated with RBD or N protein and specific IgG antibodies were detected using anti-human IgG (MH16, Sanquin).

### Statistical analysis

To investigate the differences between PBMC and BALF, effect of ICU stay and mortality on cell compositions and levels of inflammatory mediators, multiple analyses were performed wherein unsupervised clustering with OMIQ tSNE (OmiQ)) was performed. First, frequencies of immune cell subset were compared between PBMCs and BALFMCs, and soluble mediators in plasma and BALF. To aid comparability, from each patient only one sample was included in this analysis that was obtained around 14 days at the ICU and before prednisolone therapy. Statistical significance of observed differences were tested using Kruskal-Wallis (T cell differentiation and activation in PBMCs vs BALFMCs), Wilcoxon signed-rank (PD-1 expression PBMC vs BALFMC) or Mann Whitney U test (soluble mediators). PBMC populations were associated with BALFMC populations using nonlinear regression and spearman correlations without multiple-testing correction for hypothesis generation. Correlations are represented using hierarchical edge bundling plot generated using the circlize package in R as previously described.^18^. Second, to investigate the effect of ICU stay on cell compositions and inflammatory marker levels, samples were stratified in ICU stay ≤14 days and >14 days. For patients with more than one sample available in the indicated stratification, only one sample was included for the analysis (≤14 days: closest to 7 days at ICU and >14 days: closest to 21 days at ICU). Third, the association of cell composition and soluble mediators with mortality during ICU stay was investigated in both peripheral blood and BALF. In the second and third analyses statistical differences were tested using two-way ANOVA with multiple testing correction using Holm-Sidak (cell populations) or Mann-Whitney U test (soluble mediators). Statistical analysis was performed in the R statistical framework (Version 4.0.1, Vienna, Austria) or Graphad Prism v7.01 (GraphPad Software, San Diego, California, USA). Graphical presentation was performed using Graphpad Prism v7.01 (GraphPad Software), Adobe Illustrator CC v22.1 (Adobe, San Jose, California, USA), and R (Version 4.0.1).

## Author contributions

AS, EJN, LH, HJB and TvdP designed research and included patients SGB collected and analyzed patient information. AS and RL isolated samples and AS, JWD, JV, SDvA, LP performed experiments. AS, JV, SDvA, LP, YK, JJGV performed spectral flow analysis. AS, TDYR and ARS analyzed inflammatory markers. AS wrote the manuscript. TDYR, EJN, LP, ARS, SDvA, HB, TvdP and JJGV critically reviewed the manuscript. All authors reviewed and approved the manuscript.

## Acknowledgements

The authors would like to specifically thank Dorien Wouter, Selime Avci and Mariska van der Wal for their help with isolating samples and Regina de Beer, Barbara Dierdorp, Tamara Dekker and Theo Rispens for helping with measurements of soluble mediator. AS and TDY were funded in part by NACTAR (grant number 16447) of the Dutch research council (NWO) and JWD was supported by a personal grant of the NWO (VENI grant 016.186.046). On behalf of all ArtDECO consortium (Esther J. Nossent, Janwillem Duitman, Anno Saris, Heder De Vries, Lilian J. Meijboom, Lieuwe D. Bos, Siebe G. Blok, Alex R. Schuurman, Tom D.Y. Reijnders, F. Hugenholtz, Juan J. Garcia Vallejo, Hetty Bontkes, Alexander P.J. Vlaar, Joost Wiersinga, René Lutter, Tom van der Poll, Harm Jan Bogaard, Leo Heunks) and Amsterdam UMC COVID study group (S. de Bruin, A.R. Schuurman, R. Koing, M.A. van Agtmael, A.G. Algera, F.E.H.P. van Baarle, D.J.C. Bax, M. Beudel, H J Bogaard, M. Bomers, L.D.J. Bos, M. Botta, J. de Brabander, G.J. de Bree, M. Bugiani, E.B. Bulle, O. Chouchane, A.P.M. Cloherty, P.E. Elbers, L.M. Fleuren, S.E. Geerlings, B.F. Geerts, T.B.H. Geijtenbeek, A.R.J. Girbes, A. Goorhuis, M.P. Grobusch, F.M.J. Hafkamp, L.A. Hagens, J. Hamann, V. C. Harris, R. Hemke, S.M. Hermans, L.M.A. Heunks, M.W. Hollmann, J. Horn, J.W. Hovius, M.D. de Jong, N. van Mourik, J.F Nellen, F. Paulus,T.D.Y. Reijnders, E. Peters, T. van der Poll, B. preckel, J.M. Prins, s.j.raasveld, M. Schinkel, M.J. Schultz, K. Sigaloff, M.R. Smit, C. Stijnis, W. Stilma, C.E. Teunissen, P. Thoral, A.M. Tsonas, M. van der Valk, d.p.veelo, H. de Vries, M. van Vugt, D. Wouters, A.H. Zwinderman, M.C. Brouwer, W.J. Wiersinga, A.P.J. Vlaar, D. van de Beek). Funders were not in any way involved in study design or writing the manuscript.

## References

1 WHO – Coronavirus disease (COVID-19) pandemic. https://www.who.int/emergencies/diseases/novel-coronavirus-2019 (accessed Aug 18, 2020).

2 Phua J, Weng L, Ling L, et al. Intensive care management of coronavirus disease 2019 (COVID-19): challenges and recommendations. Lancet Respir Med 2020; 8: 506–17.

3 Tay MZ, Poh CM, Rénia L, MacAry PA, Ng LFP. The trinity of COVID-19: immunity, inflammation and intervention. Nat Rev Immunol 2020; 20: 363–74.

4 Krysko D V, Garg AD, Kaczmarek A, Krysko O, Agostinis P, Vandenabeele P. Immunogenic cell death and DAMPs in cancer therapy. Nat Rev Cancer 2012; 12: 860–75.

5 Merad M, Martin JC. Pathological inflammation in patients with COVID-19: a key role for monocytes and macrophages. Nat Rev Immunol 2020; 20: 355–62.

6 Vabret N, Britton GJ, Gruber C, et al. Immunology of COVID-19: Current State of the Science. Immunity 2020. DOI:10.1016/j.immuni.2020.05.002.

7 Vabret N, Britton GJ, Gruber C, et al. Immunology of COVID-19: Current State of the Science. Immunity 2020; 1. DOI:10.1016/j.immuni.2020.05.002.

8 Diao B, Wang C, Tan Y, et al. Reduction and Functional Exhaustion of T Cells in Patients With Coronavirus Disease 2019 (COVID-19). Front Immunol 2020; 11: 1–7.

9 Grifoni A, Weiskopf D, Ramirez SI, et al. Targets of T Cell Responses to SARS-CoV-2 Coronavirus in Humans with COVID-19 Disease and Unexposed Individuals. Cell 2020; 181: 1489–1501.e15.

10 Anft M, Paniskaki K, Blazquez-Navarro A, et al. COVID-19 progression is potentially driven by T cell immunopathogenesis. medRxiv 2020;: 2020.04.28.20083089.

11 Oja AE, Saris A, Ghandour CA, Kragten NAM, Boris M. Divergent SARS-CoV-2-specific T and B cell responses in severe but not mild COVID-19. medRxiv 2020. DOI:https://doi.org/10.1101/2020.06.18.159202.

12 Zhang B, Zhou X, Qiu Y, et al. Clinical characteristics of 82 death cases with COVID-19. medRxiv 2020;: 2020.02.26.20028191.

13 Wong JJM, Leong JY, Lee JH, Albani S, Yeo JG. Insights into the immuno-pathogenesis of acute respiratory distress syndrome. Ann Transl Med 2019; 7: 504.

14 McKechnie JL, Blish CA. The Innate Immune System: Fighting on the Front Lines or Fanning the Flames of COVID-19? Cell Host Microbe 2020; 27: 863–9.

15 Liao M, Liu Y, Yuan J, et al. Single-cell landscape of bronchoalveolar immune cells in patients with COVID-19. Nat Med 2020. DOI:10.1038/s41591-020-0901-9.

16 Xiong Y, Liu Y, Cao L, et al. Transcriptomic characteristics of bronchoalveolar lavage fluid and peripheral blood mononuclear cells in COVID-19 patients. Emerg Microbes Infect 2020; 9: 761–70.

17 Busse WW, Wanner A, Adams K, et al. Investigative bronchoprovocation and bronchoscopy in airway diseases. Am J Respir Crit Care Med 2005; 172: 807–16.

18 Gu Z, Gu L, Eils R, Schlesner M, Brors B. circlize implements and enhances circular visualization in R. Bioinformatics 2014; 30: 2811–2.

19 Jeannet R, Daix T, Formento R, Feuillard J, François B. Severe COVID-19 is associated with deep and sustained multifaceted cellular immunosuppression. Intensive Care Med 2020. DOI:10.1007/s00134-020-06127-x.

20 Valdez Y, Kyei SK, Poon GFT, et al. Efficient Enrichment of Functional ILC Subsets from Human PBMC by Immunomagnetic Selection. J Immunol 2018; 200.

21 Murphy K, Weaver C. Janeway’s Immunobiology, 8th edn. W.W. Norton & Company, 2011.

22 Heron M, Grutters JC, Ten Dam-Molenkamp KM, et al. Bronchoalveolar lavage cell pattern from healthy human lung. Clin Exp Immunol 2012; 167: 523–31.

23 Tolouei Semnani R, Moore V, Bennuru S, et al. Human monocyte subsets at homeostasis and their perturbation in numbers and function in filarial infection. Infect Immun 2014; 82: 4438–46.

24 Fujii T, Kadota JI, Mukae H, Kawakami K, Iida K, Kohno S. Gamma-delta T cells in BAL fluid of chronic lower respiratory tract infection. Chest 1997; 111: 1697–701.

25 Verma K, Ogonek J, Varanasi PR, et al. Human CD8+ CD57-TEMRA cells: Too young to be called ‘old’. PLoSOne 2017; 12: 1–14.

26 Romero P, Zippelius A, Kurth I, et al. Four Functionally Distinct Populations of Human Effector-Memory CD8 + T Lymphocytes. J Immunol 2007; 178: 4112–9.

27 Nguyen V, Mendelsohn A, Larrick JW. Interleukin-7 and immunosenescence. J Immunol Res 2017; 2017.

28 Jeannet R, Daix T, Formento R, Feuillard J, François B. Severe COVID-19 is associated with deep and sustained multifaceted cellular immunosuppression. Intensive Care Med 2020. DOI:10.1007/s00134-020-06127-x.

29 Martina MN, Noel S, Saxena A, Rabb H, Hamad ARA. Double Negative (DN) αβ T Cells; misperception and overdue.pdf. Immunol Cell Biol 2015; 93: 305–10.

30 Sterlin D, Mathian A, Miyara M, et al. IgA dominates the early neutralizing antibody response to SARS-CoV-2. medRxiv 2020;: 25–8.

31 Hydar Ali. Regulation of human mast cell and basophil function by anaphylatoxins C3a and C5a. Immunol Lett 2010; 128(1). DOI:doi:10.1016/j.imlet.2009.10.007.

32 likura M, Yamaguchi M, Fujisawa T, et al. Secretory IgA induces degranulation of IL-3-primed basophils. J Immunol 1998; 161: 1510–5.

33 Neumann J, Prezzemolo T, Vanderbeke L, et al. An open resource for T cell phenotype changes in COVID-19 identifies IL-10-producing regulatory T cells as characteristic of severe cases. medRxiv 2020. DOI:doi: https://doi.org/10.1101/2020.05.31.20112979.

34 Wei L, Wang W, Chen D, Xu B. Dysregulation of the immune response affects the outcome of critical COVID-19 patients. J Med Virol 2020;: 1–9.

35 St. John AL, Rathore APS, St John AL, Rathore APS. Early Insights into Immune Responses during COVID-19. J Immunol 2020;: ji2000526.

36 Mathew D, Giles JR, Baxter AE, et al. Deep immune profiling of COVID-19 patients reveals patient heterogeneity and distinct immunotypes with implications for therapeutic interventions. medRxiv 2020. DOI:10.1101/2020.05.20.106401.

