## Supplementory information for "Distinct cellular immune profiles in the airways and blood of critically ill patients with COVID-19"

### Supplemental information

#### Supplemental Figures

**Supplemental figure 1**

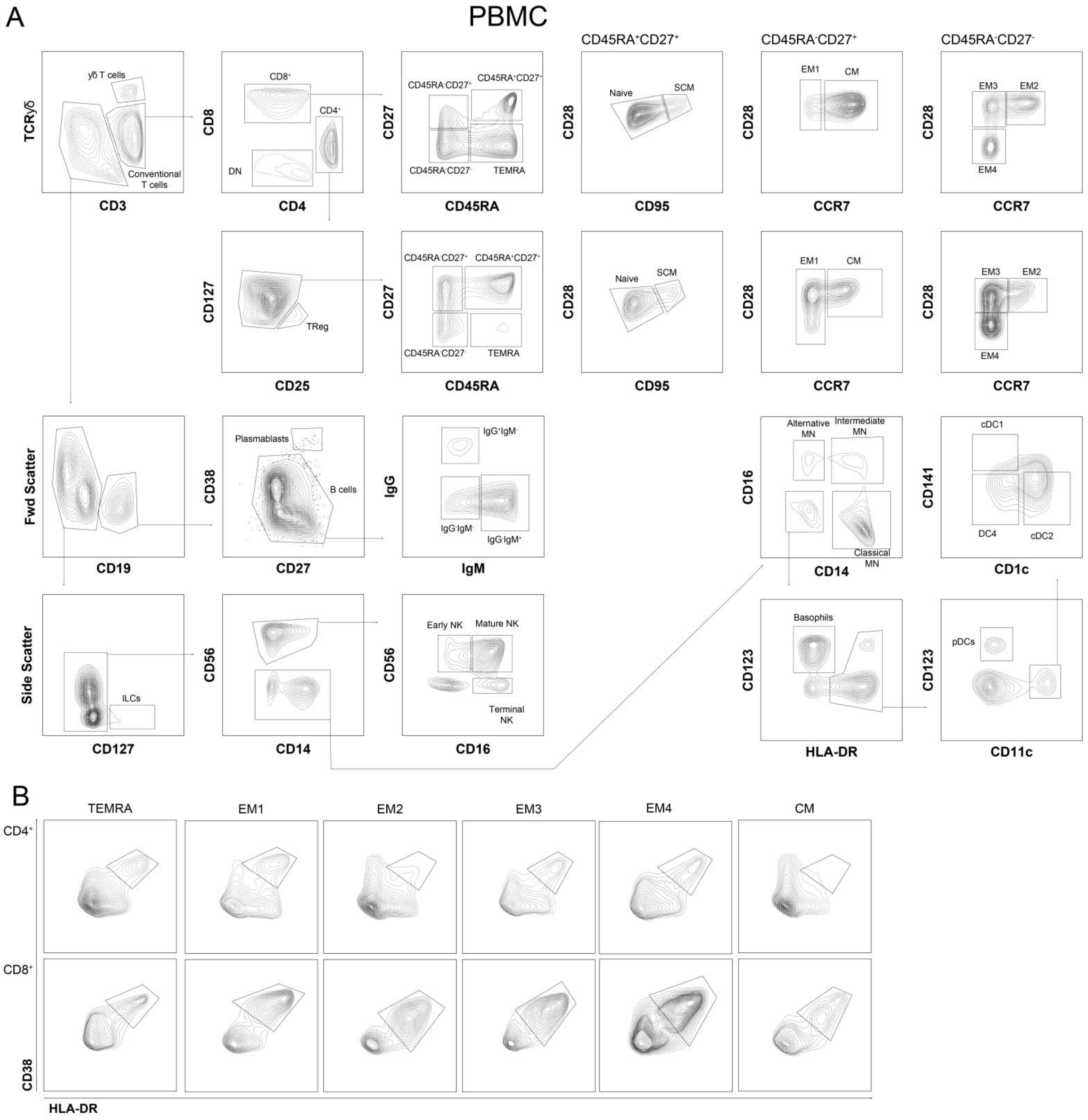


Gating strategy of cell subsets in peripheral blood mononuclear cells (PBMCs; A) and gating of T cell activation using CD38 and HLA-DR expression (B).

**Supplemental figure 2**

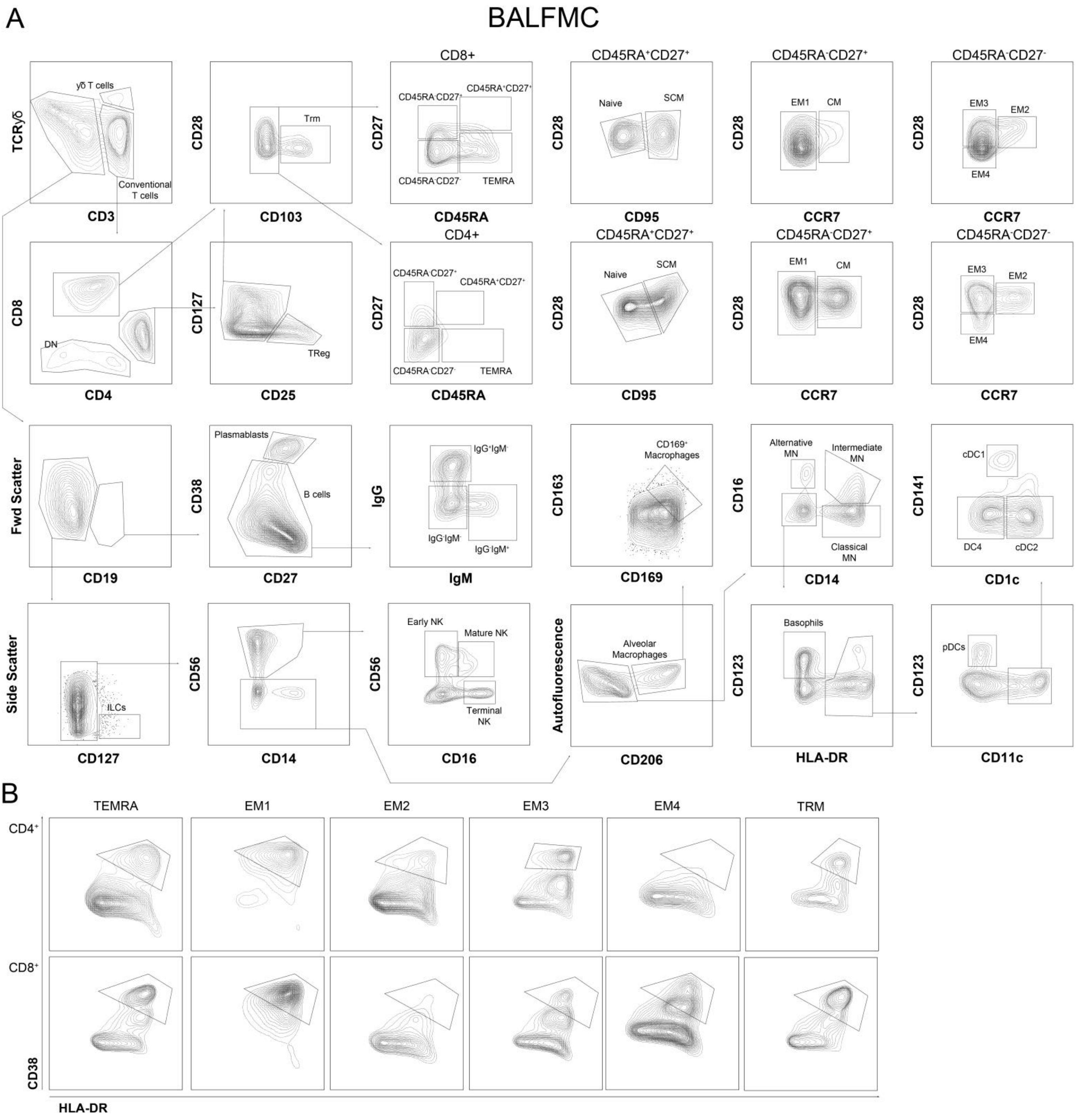


Gating strategy of cell subsets in BALF mononuclear cells (BALFMCs) (A) and of gating T cell activation using CD38 and HLA-DR expression (B).

**Supplemental figure 3**

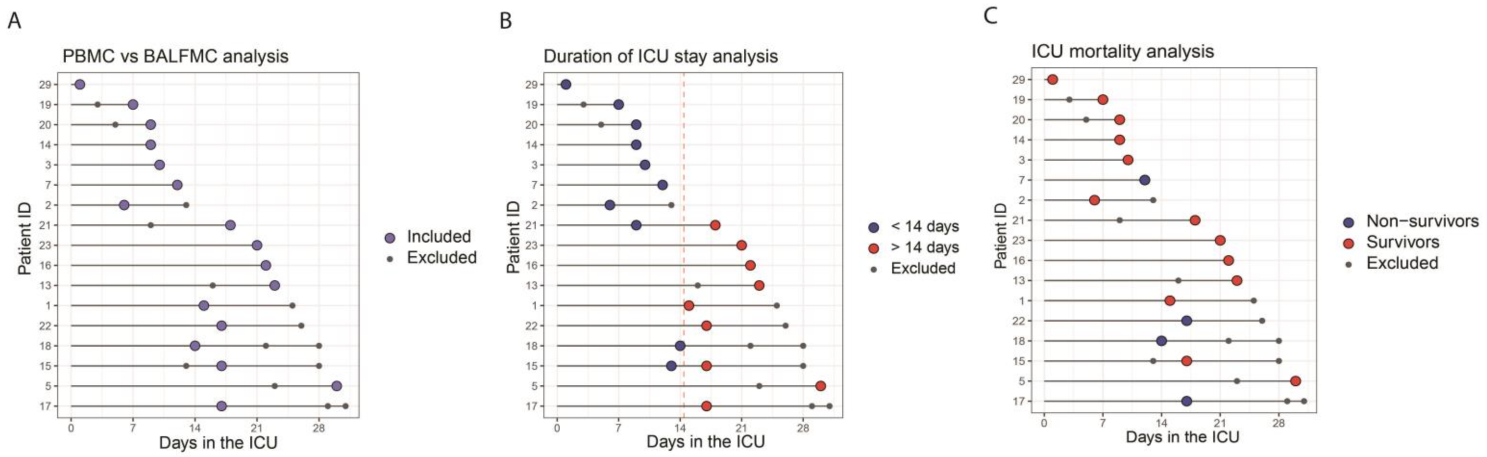


Swimmer plots displaying all patient samples that were collected of which 5 samples were excluded due to >2000 viable CD45+ cells and another 6 samples were excluded due to active prednisolone therapy. For each analysis samples that were included (red and blue circles) or excluded (small grey circles) are presented: comparing PBMCs with BALFMCs (A), duration of ICU stay (B) or ICU mortality (C).

**Supplemental figure 4**

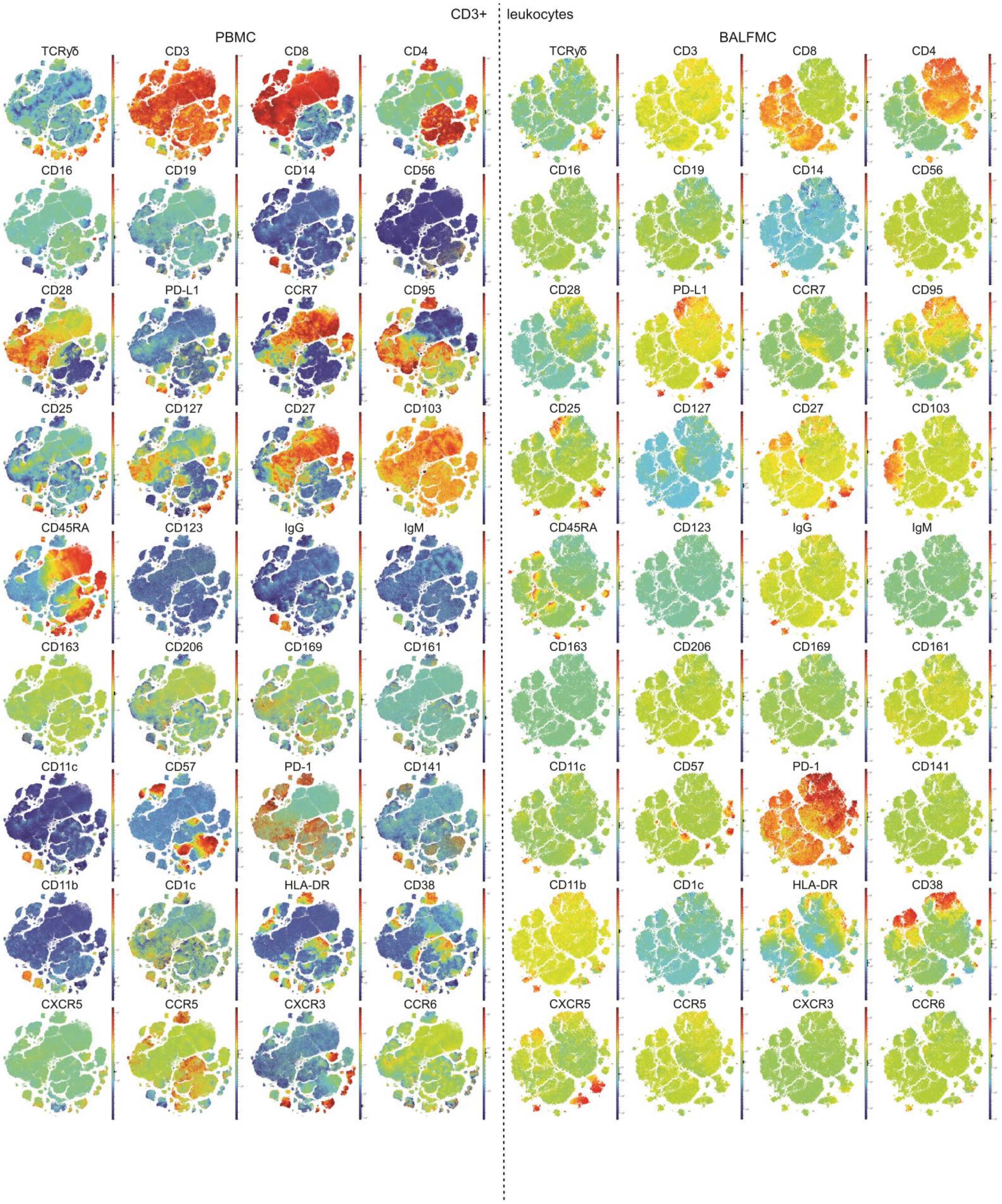


Expression patterns in viable CD45+CD3+ leukocytes of all markers that were measured. Omiq tSNE presentation of all markers individually in peripheral blood mononuclear cells (PBMCs) and BALF mononuclear cells (BALFMCs).

**Supplemental figure 5**

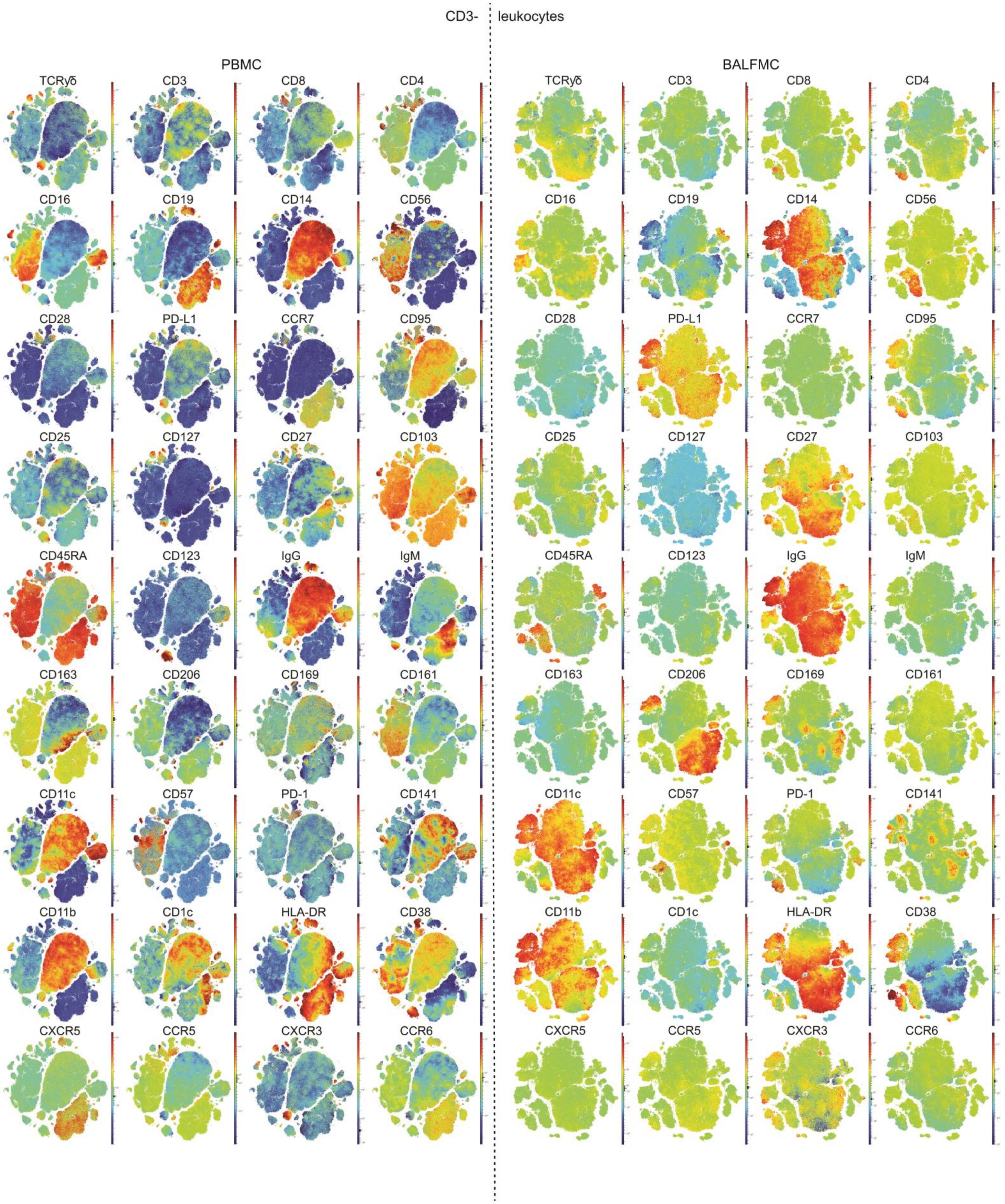


Expression patterns in viable CD45+CD3- leukocytes of all markers that were measured. Omiq tSNE presentation of all markers individually in peripheral blood mononuclear cells (PBMCs) and BALF mononuclear cells (BALFMCs).

### Supplemental tables

**Supplemental table1: overview of antibodies used in spectral flow cytometry panel**

| | Fluorochome | Catalog # | Vendor |
|---|---|---|---|
| Brilliant Stain Buffer Plus | N/A | 566385 | BD |
| LIVE/DEAD™ Fixable Blue Dead Cell Stain Kit, for UV excitation | N/A | L34962 | Thermo |
| **UV LASER** | | | |
| BUV395 Mouse Anti-Human CD45RA | BUV395 | 740315 | BD |
| BUV496 Mouse Anti-Human CD16 | BUV496 | 612944 | BD |
| BUV615 Mouse Anti-Human CD163 | BUV615 | 751355 | BD |
| BUV661 Mouse Anti-Human CD11c | BUV661 | 612967 | BD |
| BUV737 Mouse Anti-Human CD56 | BUV737 | 612766 | BD |
| BUV805 Mouse anti-Human CD169 | BUV805 | 748924 | BD |
| **VIOLET LASER** | | | |
| Brilliant Violet 421™ anti-human CD197 (CCR7) Antibody | BV421 | 353208 | Biolegend |
| CD123 Monoclonal Antibody (6H6), Super Bright 436 | Super Bright 436 | 62-1239-42 | Thermo |
| CD161 Monoclonal Antibody (HP-3G10), eFluor 450 | eFluor450 | 48-1619-42 | Thermo |
| BV480 Mouse Anti-Human CD103 | BV480 | 746742 | BD |
| Brilliant Violet 510™ anti-human CD3 Antibody | BV510 | 317332 | Biolegend |
| Brilliant Violet 570™ anti-human IgM Antibody | BV570 | 314517 | Biolegend |
| PacOrange Mouse Anti-Human CD8 | Pacific Orange | PO4958-5 | |
| BD Horizon™ BV605 Mouse Anti-Human IgG | BV605 | 563246 | BD |
| Brilliant Violet 650™ anti-human CD28 Antibody (clone CD28.2) | BV650 | 302946 | Biolegend |
| Brilliant Violet 785™ anti-human CD279 (PD-1) Antibody | BV785 | 329930 | Biolegend |
| **BLUE LASER** | | | |
| BD Horizon™ BB515 Mouse Anti-Human CD141 | BB515 | 566017 | BD |
| FITC anti-human CD57 Antibody | FITC | 359604 | Biolegend |
| Spark Blue™ 550 anti-human CD14 Antibody | SparkBlue550 | 367148 | Biolegend |
| PerCP anti-human CD45 Antibody | PerCP | 368506 | Biolegend |
| PerCP/Cyanine5.5 anti-human CD11b Antibody | PerCPCy5.5 | 301328 | Biolegend |
| TCR gamma/delta Monoclonal Antibody (B1.1), PerCP-eFluor 710 | PerCP-eF710 | 46-9959-42 | Thermo |
| **YELLOW/GREEN LASER** | | | |
| PE anti-human CD274 (B7-H1, PD-L1) Antibody | PE | 329706 | Biolegend |
| CD4 CF568 | CF568 | | Cytek |
| PE/Dazzle™ 594 anti-human 206 Antibody | PEDz594 | 321130 | Biolegend |
| PE/Cy5 anti-human CD95 (Fas) Antibody | PECy5 | 305610 | Biolegend |
| PE-Alexa Fluor 700 anti-human CD25 Antibody | PE-AF 700 | MHCD2524 | Thermo |
| **RED LASER** | | | |
| APC anti-human CD27 Antibody | APC | 356410 | Biolegend |
| Alexa Fluor® 647 anti-human CD1c Antibody | AF647 | 331510 | Biolegend |
| Spark NIR™ 685 anti-human CD19 Antibody | SparkNIR685 | 302270 | Biolegend |
| APC/Fire™ 750 anti-human HLA-DR Antibody | APCF750 | 307658 | Biolegend |
| APC-R700 Mouse Anti-Human CD127 | APC-R700 | 565185 | BD |
| CD38 APC-Fire810 | APC-Fire810 | Custom | Biolegend |

**Supplemental table 2: constitution of BALF cells before ficoll isolation**

| Date | Sample ID | Patient ID | Days at ICU | Living cells % | % CD3+ | % CD16+ | % CD56+ CD3+ | % CD19+ | % CD14++ | % CD66b+ | Unidentified | Notes on 'unidentified' |
|---|---|---|---|---|---|---|---|---|---|---|---|---|
| 15-4-2020 | 1781 | 18 | 14 | panel not yet operational | | | | | | | | |
| 16-4-2020 | 1919 | 15 | 13 | | | | | | | | | |
| 16-4-2020 | 1920 | 21 | 9 | | | | | | | | | |
| 17-4-2020 | 2012 | 14 | 9 | 97.8 | 8.1 | 4.2 | 0.6 | 1.9 | 17.9 | 62.5 | 4.9 | Partly macrophages? |
| 17-4-2020 | 2013 | 17 | 17 | 94.4 | 6.6 | 4.1 | 0.4 | 2.4 | 6.6 | 32.1 | 47.7 | Mostly debris |
| 17-4-2020 | 2014 | 2 | 6 | 38.1 | 0.2 | 5.3 | 0.1 | 1.9 | 5.5 | 84.9 | 2.1 | Mostly macrophages |
| 17-4-2020 | 2015 | 3 | 10 | 90.8 | 2.6 | 1.9 | 0.1 | 1.9 | 1.8 | 10.4 | 81.2 | Mostly debris |
| 17-4-2020 | 2022 | 1 | 15 | 94.0 | 1.0 | 1.3 | 0.1 | 1.1 | 16.9 | 63.6 | 16.0 | Mostly debris |
| 19-4-2020 | 2215 | 7 | 12 | 52.2 | 9.8 | 14.9 | 15.3 | 4.8 | 24.8 | 4.9 | 25.5 | Seem to be macrophages? |
| 19-4-2020 | 2216 | 20 | 9 | 96.9 | 12.6 | 4.0 | 0.8 | 1.7 | 22.0 | 25.9 | 33.0 | Partly macrophages |
| 19-4-2020 | 2234 | 19 | 7 | 81.5 | 24.3 | 4.1 | 2.0 | 2.4 | 20.1 | 13.4 | 33.6 | Debris and macrophages? |
| 20-4-2020 | 2326 | 22 | 17 | 98.8 | 0.0 | 3.0 | 0.0 | 2.3 | 7.6 | 65.8 | 21.4 | Seem to be macrophages? |
| 20-4-2020 | 2327 | 23 | 21 | 82.7 | 0.0 | 0.6 | 0.0 | 2.7 | 0.0 | 0.3 | 96.4 | MQs?? |
| 20-4-2020 | 2328 | 15 | 17 | 97.9 | 0.0 | 4.1 | 0.1 | 2.9 | 3.2 | 89.0 | 0.6 | - |
| 21-4-2020 | 2420 | 16 | 22 | 95.2 | 0.7 | 0.3 | 0.0 | 0.4 | 8.5 | 89.0 | 1.2 | |
| 23-4-2020 | 2668 | 13 | 23 | 96.8 | 0.7 | 1.2 | 0.1 | 1.2 | 8.7 | 78.6 | 9.4 | Partly macrophages |
| 23-4-2020 | 2678 | 5 | 30 | 99.3 | 0.3 | 0.2 | 0.0 | 0.6 | 4.6 | 84.7 | 9.6 | Partly macrophages |
| 25-4-2020 | 2901 | 21 | 18 | 96.0 | 4.8 | 2.3 | 0.3 | 1.2 | 10.9 | 77.9 | 2.7 | Partly macrophages |
| 29-4-2020 | 3266 | 17 | 29 | 96.0 | 3.1 | 13.1 | 4.5 | 6.5 | 15.5 | 3.0 | 54.4 | Seem to be macrophages? |
| 18-5-2020 | 4538 | 29 | 1 | 71.1 | 2.1 | 0.3 | 0.0 | 0.3 | 3.2 | 89.0 | 5.0 | |

**Supplemental table 3: percentages of T cell differentiation with phenotyping used during analysis**

| Tc subset | Phenotyping | PBMC | | BALF MC | |
|---|---|---|---|---|---|
| | | CD4 | CD8 | CD4 | CD8 |
| Naive | CD45RA+CD27+CD28+CD95- | 39.32% | 18.42% | 1.25% | 1.43% |
| EM1 | CD45RA-CD27+CD28+CCR7- | 4.34% | 9.19% | 0.49% | 0.79% |
| EM2 | CD45RA-CD27-CD28+CCR7+ | 6.44% | 1.86% | 24.52% | 5.18% |
| EM3 | CD45RA-CD27-CD28+CCR7- | 4.96% | 6.61% | 45.41% | 5.58% |
| EM4 | CD45RA-CD27-CD28-CCR7- | 4.75% | 9.71% | 12.88% | 39.01% |
| CM | CD45RA-CD27+CD28+CCR7+ | 24.75% | 6.75% | 0.98% | 0.19% |
| TEMRA | CD45RA-CD27-CD28+ | 4.33% | 42.04% | 2.53% | 16.62% |
| SCM | CD45RA+CD27+CD28+CD95+ | 6.71% | 5.42% | 0.08% | 0.13% |
| Treg | CD25++CD127- | 4.39% | | 8.29% | |
| Trm | CD103+CD28- | | | 3.57% | 31.07% |
